# High-throughput identification of dominant negative polypeptides in yeast

**DOI:** 10.1101/418608

**Authors:** Michael W. Dorrity, Christine Queitsch, Stanley Fields

## Abstract

Dominant negative polypeptides can act as inhibitors by binding to the wild type protein or by titrating an essential ligand. Here, we use high-throughput sequencing of DNA libraries composed of fragments of yeast genes to identify dominant negative polypeptides based on their depletion during cell growth. The method uncovers numerous inhibitory polypeptides for a protein and thus is capable of defining interacting domains with exquisite resolution, even pinpointing individual residues that contact ligands.

---

Dominant negative mutants disrupt the activity of a wild-type protein present in the same cell,^1^ allowing the detailed study of cellular processes. In contrast to the effect of more common recessive loss-of-function mutations, dominant negative inhibition occurs even when wild-type gene copies are present. The predominant mechanism of action is that a dominant negative mutant retains interaction with another molecule while lacking a critical activity. In the case of proteins that function as oligomers – estimated to represent 35% of proteins^2^ – a dominant negative polypeptide might bind to a wild-type subunit to form an inactive mixed dimer or higher order structure. In other cases, a dominant negative polypeptide may titrate out an interacting protein, a DNA site or a small molecule. For example, a dominant negative fragment of the oligomerization domain of the *E. coli lac* repressor forms nonfunctional mixed tetramers;^3^ a dominant negative Rag GTPase binds to and inhibits the mTORC1 complex;^4^ and a truncated version of the human transcription factor Stat5 that retains the DNA-binding domain saturates Stat5 DNA-binding sites.^5^ Dominant negative alleles have been linked to human disease, particularly splice variants that result in truncated proteins;^6^ truncations of human β-globin that bind heme but are unable to form functional tetramers cause dominantly inherited β- thalassemia.^7^

Inhibition of protein activity via dominant negative polypeptides has multiple advantages. Unlike RNAi approaches which require degradation of the targeted protein before an effect is observed, dominant negative polypeptides can act almost immediately. Unlike CRISPR/guide RNA approaches that permanently alter a gene, dominant negative polypeptides can be conditionally induced and expressed at variable levels,^1^ features useful for inhibiting essential genes. Furthermore, because dominant negative polypeptides bind their targets based on protein structural features, not nucleic acid sequence specificity, they have the potential to inhibit individual functions of a protein. In addition, binding to structural features allows a single dominant negative polypeptide to inhibit multiple proteins that possess a common domain. Despite their utility, dominant negative alleles are typically discovered only at small scale in the context of broader mutant screens.^8^ Larger-scale attempts have been made in the yeast *Saccharomyces cerevisiae* to identify polypeptide fragments that, when overexpressed, produce dominant negative effects, but these studies were limited in their sensitivity and their capacity to track potential inhibitory fragments.^9–12^

We used high-throughput sequencing of DNA libraries encoding *S. cerevisiae* polypeptide fragments to identify thousands of dominant negative inhibitors under different selective conditions. Comparing the frequencies of fragments present in yeast cells before and after induction, we identified fragments associated with decreased cellular fitness. We initially tested the method with individual proteins known to oligomerize. Yeast cells require homodimers of orotidine-5’-phosphate decarboxylase (Ura3 protein) to synthesize pyrimidines and thus to grow in media lacking uracil. We fragmented the *URA3* gene to ∼200 bp, and cloned >7,000 fragments into a low-copy vector with a galactose-inducible promoter, which when induced should produce 25- to 100-fold more fragment than the full-length Ura3 expressed from its native promoter.^13,14^ Triplicate cultures transformed with this library were induced with galactose and grown 24 generations in media lacking uracil (Fig. 1A). Cells were collected before induction and 48 hours later. We sequenced inserts to track the frequencies of each fragment before and after selection, allowing us to calculate a depletion value for each fragment based on its change in frequency. Values ranged from highly depleted (2-6% of starting frequency) to neutral (similar to starting frequency) (Fig. 1A, lower panel).

**Figure 1.**
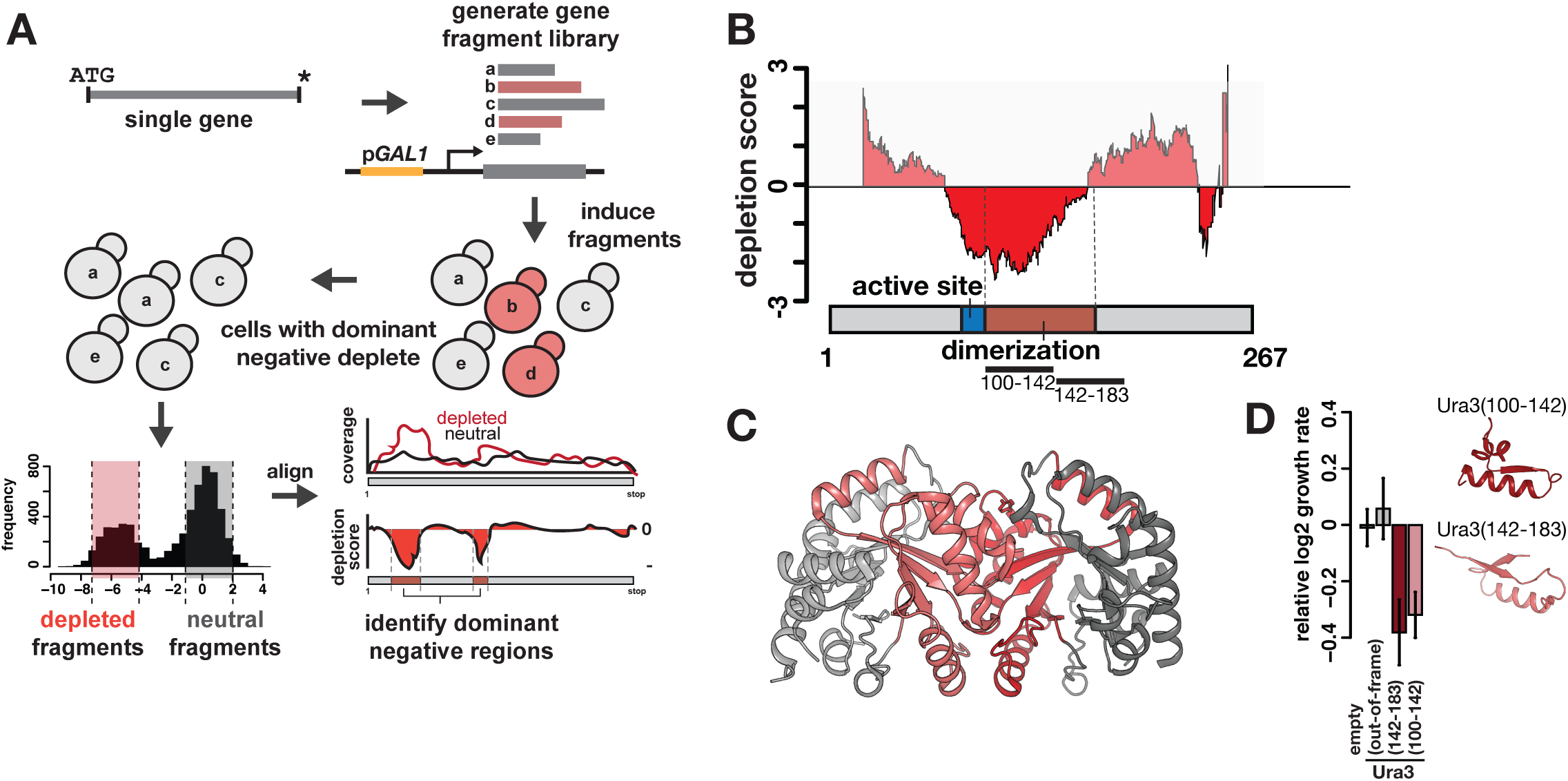
Dominant negative polypeptides can be identified in high-throughput and correspond to known protein domains. (A) A schematic of the experimental and computational pipeline to identify dominant negative polypeptides in high-throughput. (B) Dominant negative polypeptides are enriched, relative to neutral polypeptides, in a central region of the Ura3 protein. Negative depletion scores indicate regions with excess dominant negative fragments. Positions on either end of Ura3 with no computed depletion score had fewer than 15 mapped fragments. A model of the Ura3 protein is shown below, with active site and dimerization domain highlighted. (C) The dominant negative region identified in (B) was mapped onto the crystal structure of the Ura3 homodimer; this region covers nearly all residues in the homodimer interface. Two individual fragments (amino acids 100-142 and amino acids 142-183) selected for validation are shown. (D) Barplots showing individual growth rates for yeast cells containing a wild-type copy of the *URA3* gene and one of three fragments (amino acids 10-43 out of frame, amino acids 100-142, amino acids 142-183). Growth rates are normalized to a strain growing without fragment expression. Error bars represent standard error of the mean.

While most (981/1318, 74.4%) in-frame fragments had little to no effect, 337 (25.6%) showed dominant negative activity, depleting to less than 1/16 their starting frequency. We aligned these to the *URA3* gene, controlling for coverage biases by also aligning equivalent numbers of inframe fragments with neutral depletion scores and out-of-frame fragments, to establish a perposition depletion score. Depleted fragments and per-position depletion scores correlated (*r*^*2*^=0.94, *r*^*2*^=0.9) across replicates (Suppl. Fig. 1). The region most depleted encompassed amino acids 70 to 160 (Fig. 1B), a region that corresponds to the Ura3 dimer interface (Fig. 1C). This region also contains active site residues required to convert orotidine-5’-phosphate into uridine monophosphate (Fig. 1B, below). The most depleted residue within this region is amino acid 100 (Fig. 1B), a threonine that makes a critical substrate contact,^15^ and the smallest inhibitory polypeptides (<30 amino acids) included residues near the active site (Suppl. Fig. 2). The inhibitory region (Fig. 1C, red) contains four α-helices and two β-strands. Two fragments from this region reduced growth in media lacking uracil by 25%, while an out-of-frame control fragment had no effect (Fig. 1D). As this approach does not exclude the identification of depleted fragments with non-specific effects, such as forming toxic aggregates or disrupting membranes, we tested for growth defects due to the putative Ura3 dominant negative fragments in media containing uracil, where the Ura3 protein is not essential. We found no growth defects in this condition (Suppl. Fig. 5).

We carried out a similar experiment with the transcription factor Hsf1, which contains three conserved domains: a DNA-binding domain, a coiled-coil helix required for homo-trimerization, and a putative repressive autoregulatory helix. Hsf1 activates genes required for basal growth at 30°C and the heat-shock response at 37°C, and is essential at both temperatures. We generated a library of >12,000 fragments of *HSF1*, transformed yeast, induced expression and grew cells at either 30°C or 37°C. Most fragments showed no change in depletion across the selection at either temperature, but 377 (19.8%) in-frame fragments were depleted (1/16 of starting frequency or less) under basal condition, and 436 (23%) under heat shock. These depleted fragment sets largely overlapped (319/494, 64.6%), suggesting that most dominant negative Hsf1 polypeptides act independent of temperature. Alignments of depleted and control fragment sets reveal three regions with dominant negative effects (Fig. 2A), corresponding to the conserved domains.

**Figure 2.**
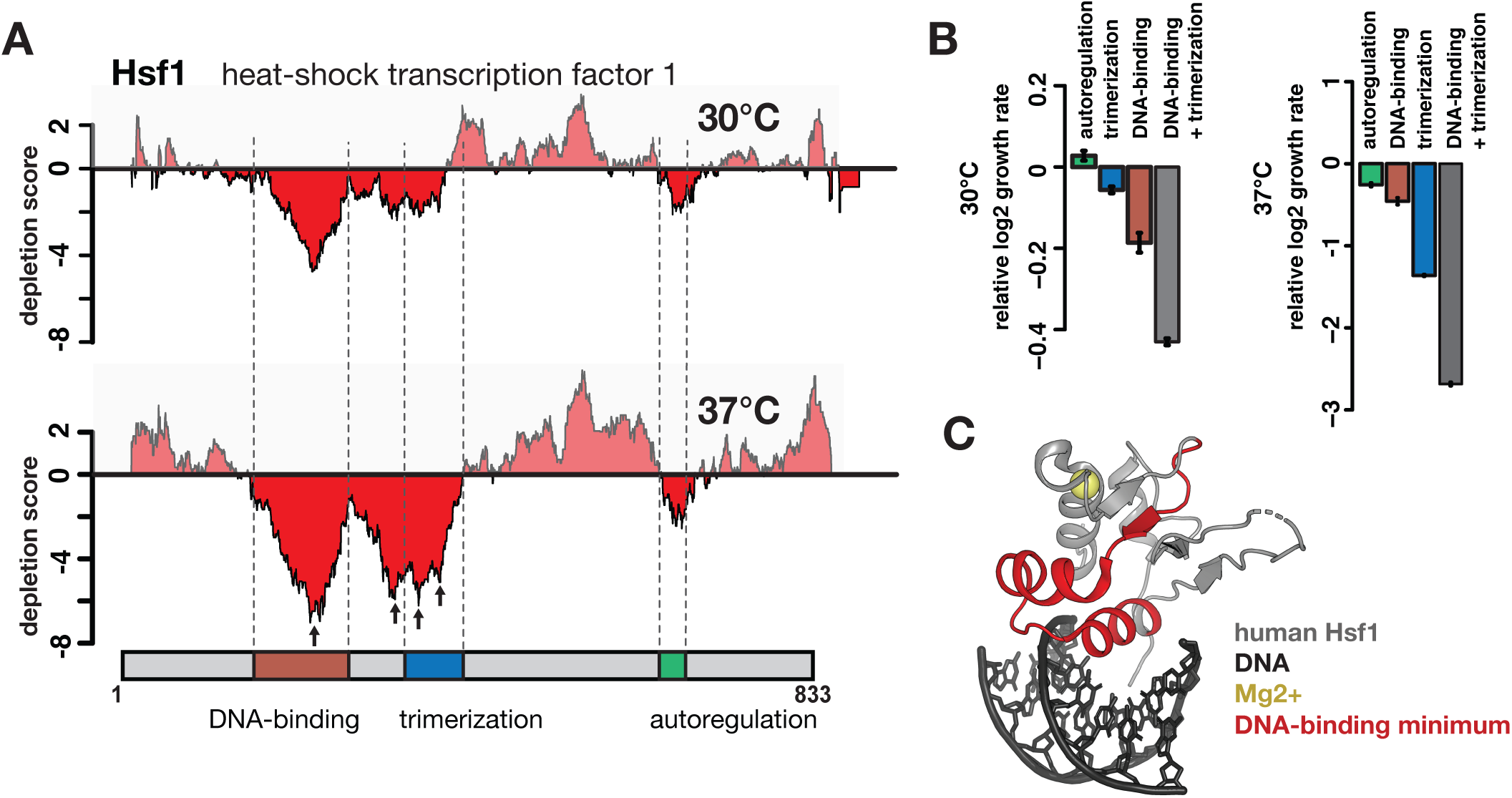
Identifcation of conditional dominant negative polypeptides in the essential heat-shock transcription factor Hsf1. (A) Plots showing depletion scores as in Fig. 1B for dominant negative identification in Hsf1. Depletion scores for cells grown in basal conditions are shown in the top panel, and scores for cells grown under heat-shock are shown in the lower panel. Dominant negative regions overlapped functional domains of Hsf1 (indicated with dashed lines). Local minima are indicated with arrows. (B) Individual growth rates for the regions identified in (A) reveal increased dominant negative activity of a trimerization domain fragment under heat-shock conditions, as well as a role for the short autoregulatory domain. Error bars represent standard error of the mean. (C) Local minimum in the DNA-binding domain from (A) is mapped onto the structure of the human Hsf1 DNA-binding domain.

To validate these effects, we measured basal and heat-shock growth rates of cells expressing one fragment from each domain. The DNA-binding domain fragment was the most deleterious, at 30°C and 37°C (Fig. 2B). In contrast, the trimerization domain fragment reduced growth at 30°C by only 4%, but under heat-shock by 56% (Fig. 2B), consistent with previous dominant negative Hsf1 mutants^16^ and the production of nonfunctional trimers with wild-type Hsf1 subunits. The fragment corresponding to the autoregulatory domain in the human Hsf1 protein had no effect at 30°C, but reduced growth at 37°C by 15% (Fig. 2B), suggesting that this domain has conserved activity in *S. cerevisiae*, despite the assertion that it is nonfunctional in this organism.^17–19^ We mapped the minima of the dominant negative regions onto the crystal structure. The minimum in the DNA-binding domain (Fig. 2A) centers on the recognition helix that contacts the major groove of DNA (Fig. 2C). Three other minima include two regions in the linker that precedes the predicted trimerization helix, implicated in promotion of DNA binding,^20^ and a region spanning the first two heptad repeats of this helix (Fig. 2A). Thus, the dominant negative strategy revealed new features of the extensively-studied Hsf1, such as the activity of an autoregulatory domain in yeast.

To test this approach across many genes at once, we transformed yeast with a library of >172,000 yeast cDNA fragments and imposed a growth selection over 48 hours to obtain depletion scores. Of 6,713 annotated yeast genes, 3,311 had at least one in-frame fragment with adequate sequencing depth (see Methods). Of these 3,311, 1,866 had at least one strongly (1/250 of starting frequency or less) depleted in-frame fragment, representing possible dominant negative inhibitors.

We next sought to identify gene regions enriched for dominant negative fragments. We identified the 20,000 most depleted in-frame fragments, along with an equal number of out-of-frame and neutral control fragments, and mapped these back to individual genes. We filtered this set to genes with at least 5 in-frame fragments, identifying 456 genes (corresponding to 6,327 fragments). We filtered for genes with at least 4 overlapping fragments and genes with more inframe than out-of-frame fragments (see Methods), leaving a set of 157 genes for which a dominant negative region could be readily identified (Suppl. Fig. 3; Suppl. Data 1). These 157 genes were enriched for components of the ribosome and translation overall, consistent with the essential nature and the abundance of these proteins as encoded in a cDNA library (Fig. 3A). Fourteen ribosomal proteins contained regions with overlapping dominant negative fragments (Suppl. Fig. 4), eleven derived from the cytoplasmic ribosome and three from the mitochondrial ribosome. Most yeast ribosomal proteins are encoded by highly similar paralog pairs. The eleven genes include two with dominant negative regions from both paralogs and two from only one member of the paralog pair. Ribosomal proteins are monomeric, indicating that these dominant negative polypeptides are not interfering with homo-oligomerization and may instead be inhibiting ribosome assembly. The set of dominant negatives derived from ribosomal proteins reveals that large complexes can be inhibited by a fragment from a single protein in the complex (Suppl. Fig 4). Seven of the cytoplasmic ribosomal proteins could be mapped onto the cryo-EM structure^21^ of the eukaryotic translating ribosome (Fig. 3A; Suppl. Fig. 4), showing regions of this complex sensitive to dominant negative inhibition.

**Figure 3.**
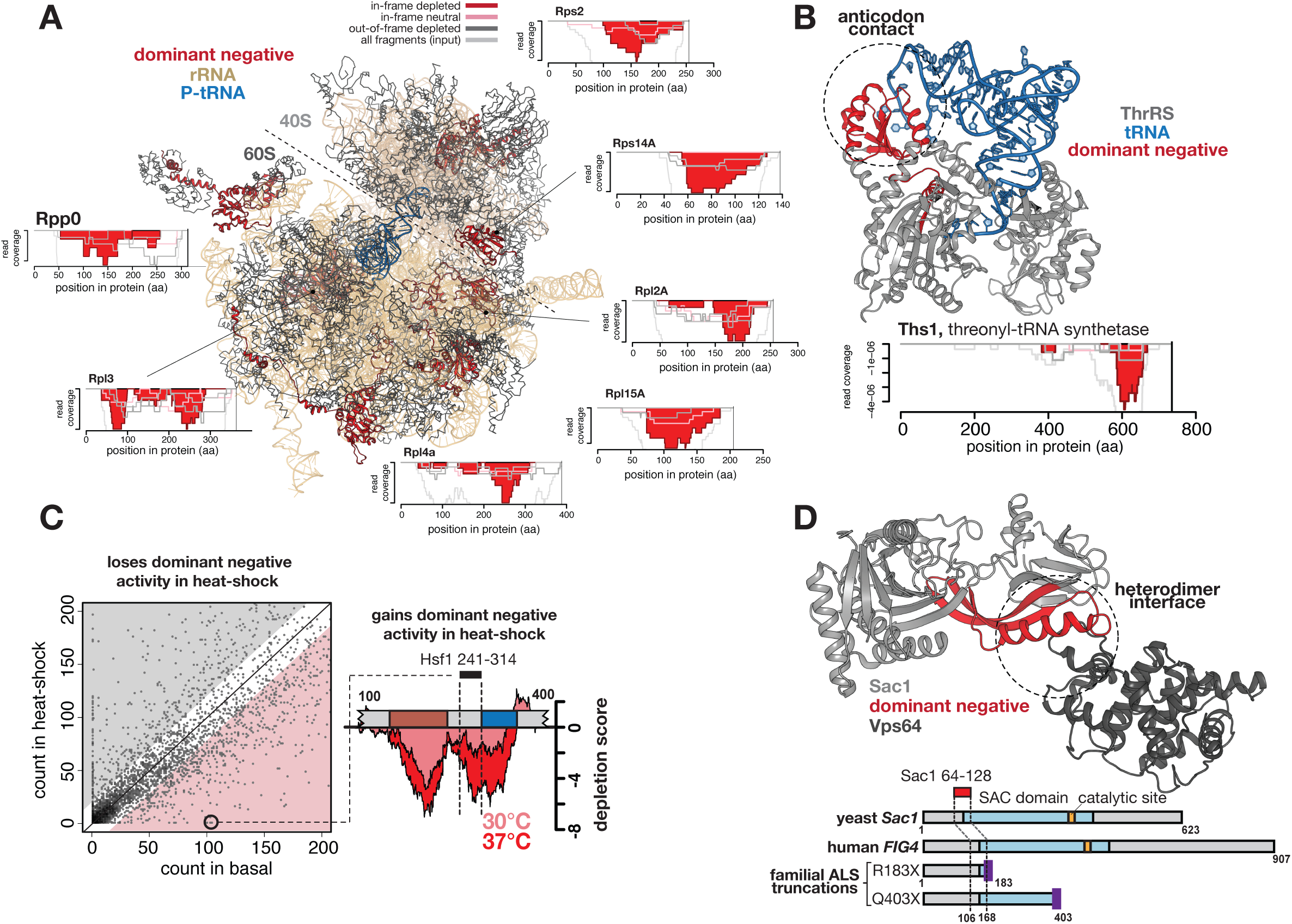
Dominant negative inhibitors identified in full-genome screen. (A) Ribosomal proteins were mapped onto the cryo-EM structure of the translating ribosome, with the small subunit (40S) shown in gray, the large subunit (60S) in darker gray, the 18S and 25S rRNAs in gold, the P-site tRNA in blue, and proteins with dominant negative polypeptides in red. Around the structure are depletion plots that show domains in these proteins with overlapping dominant negative polypeptides. Y-axis of each depletion plot is coverage frequency normalized to total coverage (see Methods; Suppl. Data 1). (B) The dominant negative region identified in the threonyl-tRNA synthetase (red) was mapped onto the crystal structure (gray) in complex with its cognate tRNA (blue). The dominant negative domain (red) corresponds to the anticodon binding region of the tRNA synthetase. (C) The frequency of each fragment in basal (x-axis) and heat-shock (y-axis) conditions shows that many fragments lose their activity under heat-shock, and a subset gain dominant negative activity under heat-shock. The fragment with the greatest heat-shock-specific dominant negative activity derived from the linker region between the DNA-binding domain and trimerization domain of Hsf1 (inset from Fig. 2A). (D) A temperature-sensitive dominant negative fragment (red) derived from the yeast Sac1 gene is highlighted at a heterodimer interface in the crystal structure of the Sac1-Vps64 complex. Below is a schematic representation of the yeast Sac1 protein and its human homolog *FIG4*. The SAC domain is shown in light blue, and the catalytic site in orange. Two truncation variants of human *FIG4* associated with dominant ALS are shown below, and the conserved position of the dominant negative fragment identified in yeast is shown with dashed lines.

We observed hits in other proteins necessary for translation, including the domain of tRNA synthetases that binds the anticodon (Fig. 3B, Suppl. Fig. 5), as shown for a previous tRNA synthetase dominant negative.^22^ We found dominant negative regions in genes for the cell cycle (*PCL5, SRM1, RME1, CDC34*); amino acid metabolism (*LYS14, CDC34, HIS1, THR1, ADE3, ARO1, ARO8*); tRNA processing (*STP4, STP1, SRM1, SOL3, RPN2, SEN1*); stress response (*SSA1, TPS1, GPD1, SSA4, HSP12, SSC1, UTH1, SSA2, HSP60, SSE1*); cytoskeleton (*ARC15, LSB5, RVS167*); and cell wall maintenance (*CCW12, CWP2, UTH1*), suggesting that this method is broadly applicable in targeting cellular processes.

We tested fragments from three biosynthetic genes, *ADE3* (for adenine), *THR1* (for threonine), and *ARO1* (for aromatic amino acids), for non-specific toxicity. Identification of these fragments in the large-scale selection is consistent with the growth defects of null mutants in these genes even in complete media.^23^ We asked whether growth defects due to these fragments were magnified when the relevant nutrient was not provided. Overexpression of the most depleted fragment from Thr1 (300-350) further reduced growth rate by 17% in media lacking threonine, and from Ade3 (882-933) by a further 16% in media lacking adenine (Suppl. Fig. 6). The Thr1 fragment had no effect in media lacking adenine, nor did the Ade3 fragment in media lacking threonine, suggesting that these fragments are specific to their targets (Suppl. Fig. 6). A fragment from Aro1 (1491-1551) reduced growth in media lacking phenylalanine, but so did fragments from Thr1 and Ade3, suggesting that growth in this media cannot provide evidence for specificity. Overall, these results suggest that non-specific toxicity is not a primary signal in the data.

To identify condition-specific inhibitors, we applied a heat stress during the growth of cells expressing the cDNA library. The large majority (97.8%) of dominant negative polypeptides that depleted in basal conditions also depleted under heat shock (Fig. 3C). We found 508 dominant negative polypeptides with depletion to <1/16 starting frequency only in heat shock, with the most extreme 26 of these missing under heat shock (Fig. 3C). The most strongly depleted fragment corresponded to the linker region preceding the trimerization domain of Hsf1, which we had identified as having temperature-specific dominant negative activity (Fig. 3C, right side inset from Fig. 2A).

We found a dominant negative fragment derived from the phosphatase Sac1 (64-128) that depleted to 2% of its basal level and that corresponds to an interface with its interacting partner Vps64 (Figure 3D). Protein truncations of the human homolog of Sac1, *FIG4*, occur in heterozygous form and are associated with familial amyotrophic lateral sclerosis,^24^ suggesting that dominant negative activity may contribute to this disease. That polypeptide fragments expressed in both basal and heat-shock conditions inhibited growth only at high temperature suggests that their mode of action is revealed under stress. We also found about twice as many fragments (1,248) that showed the opposite trend, with strong dominant negative activity (depleted to <1/16 starting frequency, 126 fully dropped out) only in basal growth conditions. The high temperature may affect the availability of the target or the binding interaction itself.

The method developed here can be applied to specific genes or entire libraries from any organism with suitable genetic tools. The timing, magnitude, and spatial pattern of inhibitor activity can be precisely controlled using inducible expression. The method can be readily adapted to screens and selections besides viability, including ones that can be performed under multiple conditions to obtain condition-specific inhibitors. The resulting data should allow the definition of functional domains corresponding to interaction surfaces, particularly domains that mediate homo-oligomerization or protein-protein interaction, although the mere identification of a dominant negative polypeptide does not distinguish its mechanism of action.

Remarkably, the method is capable of pinpointing residues corresponding to local minima that may be of special relevance, such as contacting RNA, DNA or small molecules. These exceptional residues can serve as anchor points to select small fragments with dominant negative activity, which may be useful in the design of therapeutics.^25,26^ Some proteins have multiple minima that are distant in the primary sequence but map to nearby sites in the three-dimensional structure. In 3’-phosphoglycerate kinase (Pgk1), these minima correspond to features below the level of the protein domain, but not to known ligand-binding sites or interaction interfaces (Suppl. Fig. 7); the inhibition may relate to protein folding. Although the dominant negatives we identified are from yeast, the homology of many of them to proteins from human cells suggests that similar small fragments of mammalian proteins can be found. Given the diversity of possible selection schemes, this method can address such biological questions as protein-based species-incompatibilities, fine-tuning of biological circuits at the protein level, and condition-specific and cell-type-specific protein interactions and structures, in addition to providing highly precise and rapidly acting reagents for functional genomics.

## Methods

### Fragment library construction and transformation

For individual gene libraries, the *URA3* and *HSF1* genes were amplified via PCR, and fragmented to an average length of 200 bp using Illumina Nextera transposition. To generate a cDNA-derived fragment library, RNA was isolated from yeast cells via acid-phenol extraction, and polyA-specific cDNA was synthesized and transposed to obtain gene fragments. Transposed fragments were amplified via PCR and cloned into a low copy, galactose-inducible expression vector. Because each fragment contained a 5’ adaptor sequence (AGATGTGTATAAGAGACAG) and 3’ adaptor sequence (CTGTCTCTTATACACATCT) resulting from the transposition, the vector was designed to translate through these sequences such that each in-frame fragment would be translated as **MKDVYKRQ**-NNN-**LSLIHILTD**\*\*. Electromax *E. coli* cells were transformed with the library assemblies and used to amplify a large pool of plasmid for subsequent yeast transformation. Yeast cells (W303 strain) were transformed with fragment libraries such that each cell received a low-copy plasmid carrying a *TRP1* gene for plasmid selection.

### Cell growth selections

For the *URA3* experiment, yeast cells containing the *URA3* fragment library were grown without selection for Ura3 function in synthetic complete medium lacking tryptophan (SC-TRP, for plasmid selection only). After 2 hours of pre-induction with galactose, these cells were washed to remove excess media and transferred to induction media lacking uracil (SC-TRP-URA, +galactose) for the selective growth condition. After reaching log phase, the cells were back-diluted and grown again in fresh SC-TRP-URA, +galactose for another round of growth selection. The *HSF1* experiment was conducted in a similar fashion, except that after the 2 hour pre-induction, the culture was split such that one population continued growth at 30°C (SC-TRP, +galactose), and the second population was shifted to 37°C (SC-TRP,+galactose). Each culture was back-diluted after reaching log phase to undergo additional growth selection. Growth selections for the cDNA experiment were conducted as for the *HSF1* experiment. In the growth selections, care was taken to transfer enough cells for at least a 1000-fold coverage of the initial fragment library size.

### High-throughput sequencing and analysis of polypeptide fragments

Sequencing was carried out on Illumina’s NextSeq platform. DNA sequencing libraries were prepared from plasmids extracted from yeast populations (Yeast Plasmid Miniprep II, Zymo Research, Irvine, CA) before and after selection, as well as from the initial plasmid input. These plasmids were used as template for PCR amplification (< 15 cycles) that added sequencing adaptors, as well as 8 bp sample indexes. Paired-end reads spanning the fragment library were generated at a median of 1 million reads per sample for individual gene fragment libraries, and between 15-50 million reads per sample for the larger cDNA fragment libraries. The position and orientation of each fragment was determined by aligning each set of reads to the respective gene of origin, or to the full set of coding ORFs (SGD). Reads were aligned using Bowtie2. The frequency of each fragment was determined by counting unique fragments with identical gene of origin, DNA start site, and DNA stop site, and orientation (ex. HSF1_202_431_+). Each fragment was assigned a reading frame based on its start position. Fragments with less than five reads were filtered from analysis steps due to inadequate coverage. Dominant negative regions of individual gene selections for Ura3 and Hsf1 were determined by computing per-position fragment coverage within each gene and taking the ratio of in-frame depleted fragments to out-of-frame depleted fragments, i.e. a depletion score (-log2[coverage_in-frame depleted_ / coverage_out-of-frame depleted_]). Dominant negative regions of genes from the cDNA library were determined by (1) filtering out genes with fewer than 5 in-frame fragments; (2) filtering out genes with fewer than 4 overlapping in-frame fragments; (3) filtering out genes whose number of overlapping in-frame fragments was less than 2-fold higher than the number of overlapping out-of-frame fragments.

## Acknowledgments

We thank Robert Waterston and Evan Eichler for comments on the manuscript. The work was supported by NIH grant GM114166 to C.Q. and S.F. and NIH grant P41 GM103533. During most of this work, S.F. was an investigator of the Howard Hughes Medical Institute.

## Author Contributions

M.W.D., C.Q., and S.F. conceived and interpreted experiments. M.W.D., C.Q., and S.F. wrote the manuscript. M.W.D. conducted experiments and data analysis.

## Data Availability

High-throughput sequencing reads have been submitted to NCBI SRA, awaiting accession number. Positional enrichment scores for each gene tested are provided in source data files.

---

**Suppl. Fig. 1.**
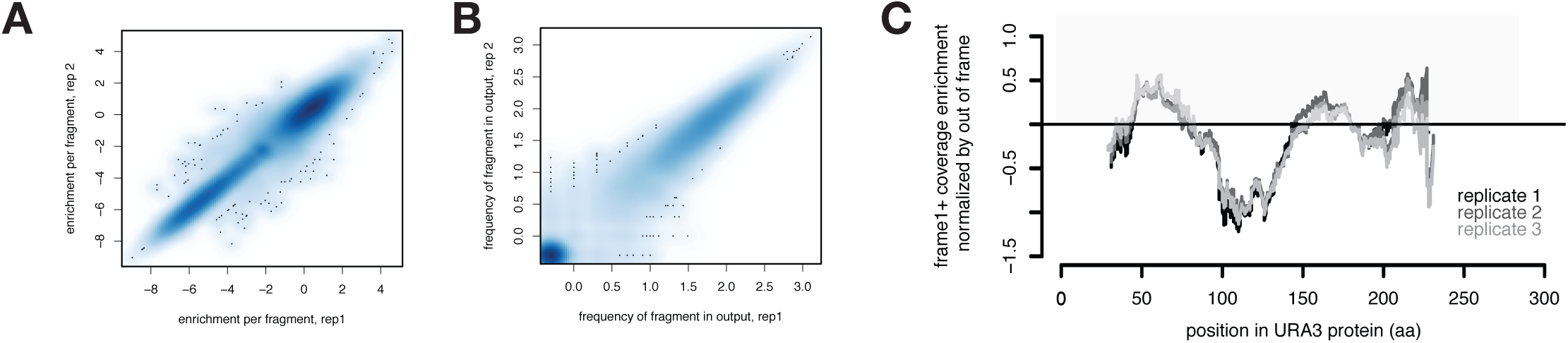
Reproducibility of individual fragment depletions and coverage values across Ura3 protein. (A) Scatterplot showing the enrichment scores (log2[frequency*out*/frequency*in*]) of each fragment of Ura3 across two biological replicates. Darker shades of blue indicate a higher density of points. (B) Scatterplot showing frequencies of each Ura3 fragment in the output cell populations in two replicates. The strong density of points in the lower left corner of the plot indicate fragments that fully dropped out during the selection in media lacking uracil. (C) Overlaid fragment coverage plots (as in Fig. 2B) generated from each of three replicates (shades of gray).

**Suppl. Fig. 2.**
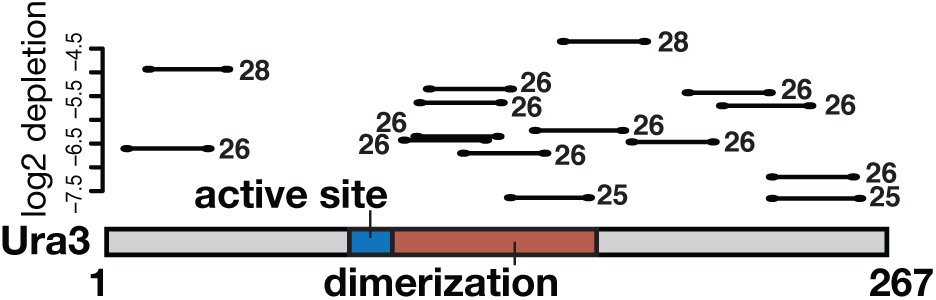
The smallest dominant negative peptides from Ura3. Plot showing the fifteen smallest and most depleted fragments from Ura3; y-axis shows the fragment enrichment score and x-axis indicates position in the Ura3 protein. The size (in amino acids) is shown beside each fragment, but all fragments contain peptide adaptors (see Methods), such that the full translated products range from 39-42 amino acids.

**Suppl. Fig. 3.**
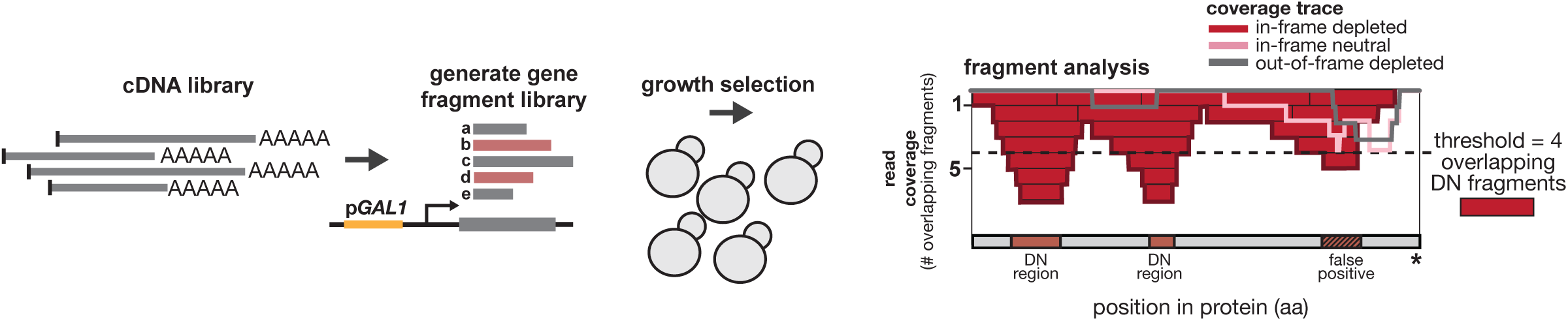
Scheme for full-genome dominant negative fragment identification. A yeast cDNA library generated from cells growing at 30°C was used to generate a library of fragments cloned into a vector with galactose-inducible expression (as in Fig. 1A). After induced yeast cells containing this library have grown overnight, fragments from the output population are sequenced along with fragments from the pre-induction population to produce enrichment scores as in Fig. 1A. After identifying the reading frame and functional effect (depleted or neutral) of each fragment from the cDNA library, these fragments were aligned back to yeast genes. Fragment pileups from the in-frame depleted (red) were compared to pileups generated from two control sets: (1) in-frame neutral fragments (pink) and (2) out-of-frame depleted fragments. Genes with dominant negative regions were identified having a minimum of four (dashed line) overlapping in-frame depleted fragments, and as having regions where coverage of the control sets do not exceed the coverage of the in-frame depleted set (example in C-terminal pileup region above).

**Suppl. Fig. 4.**
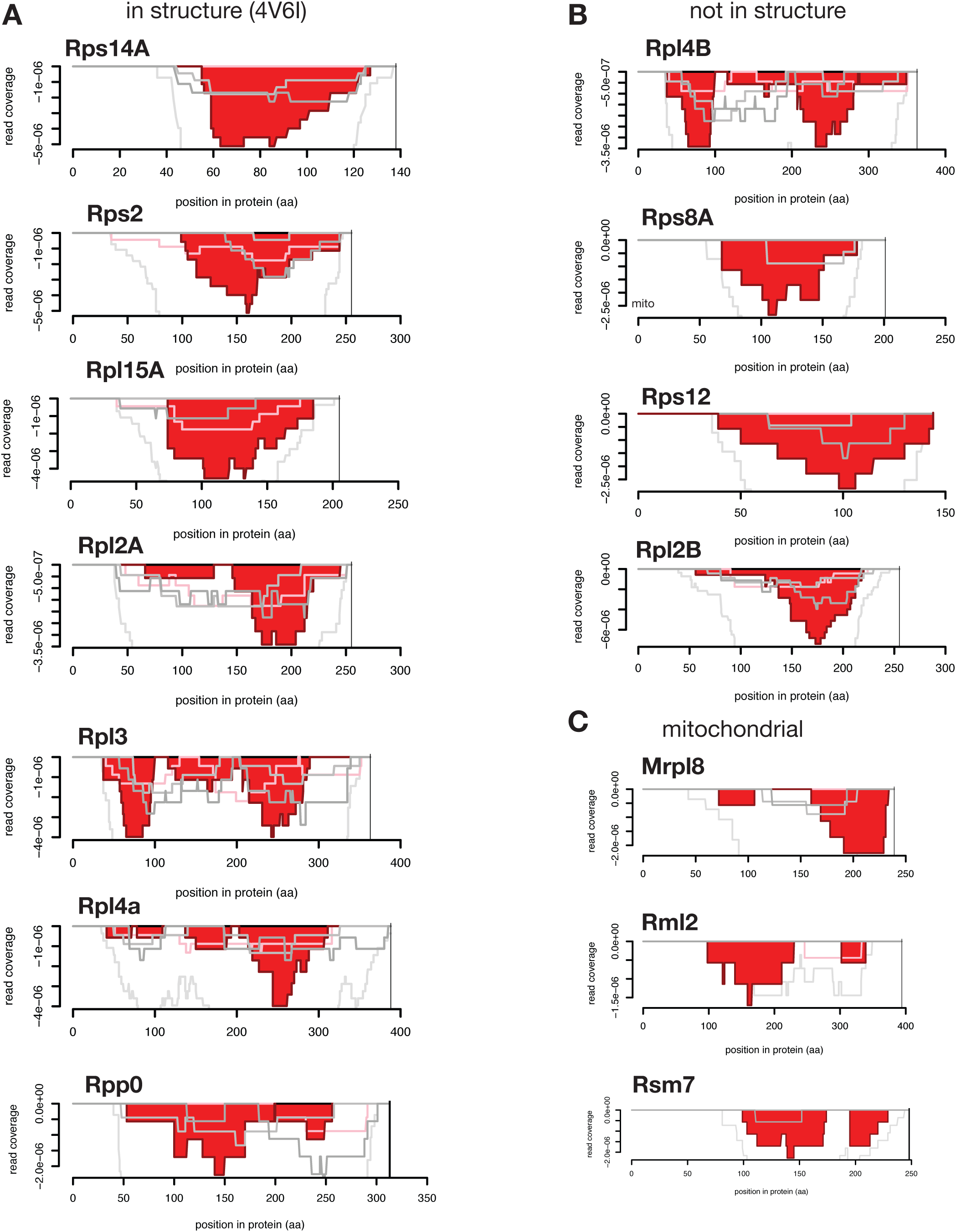
Dominant negative coverage in protein components of the yeast ribosome. Coverage charts (as in Fig. 3A) are shown for all ribosomal proteins in the yeast cDNA library screen identified to contain dominant negative regions. (A) Those present in the cryo-EM structure of the translating yeast ribosome (4V6I) are shown; (B) those absent in the structure or (C) that derive from the mitochondrial ribosome are also shown.

**Suppl. Fig. 5.**
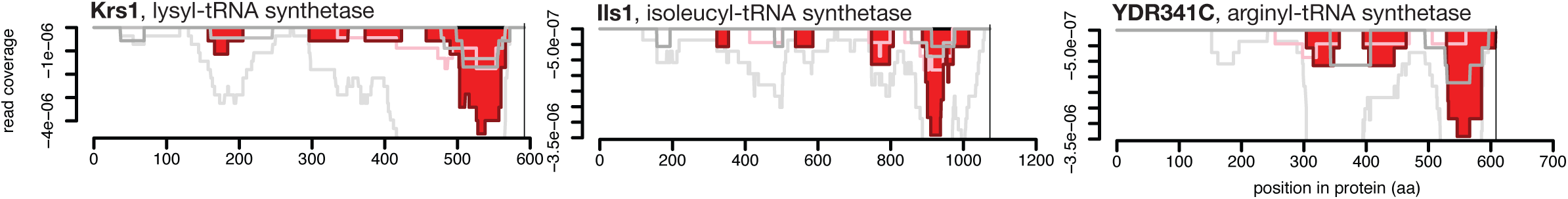
Several tRNA synthetases show overlapping dominant negative fragments in the C terminus. Coverage charts (as in Fig. 3B) are shown for all tRNA synthetase proteins in the yeast cDNA library screen identified to contain dominant negative regions. For these examples and the threonyl-tRNA synthetase shown in Fig. 3B, the dominant negative regions were found in the C-terminal end of the protein.

**Suppl. Fig. 6.**
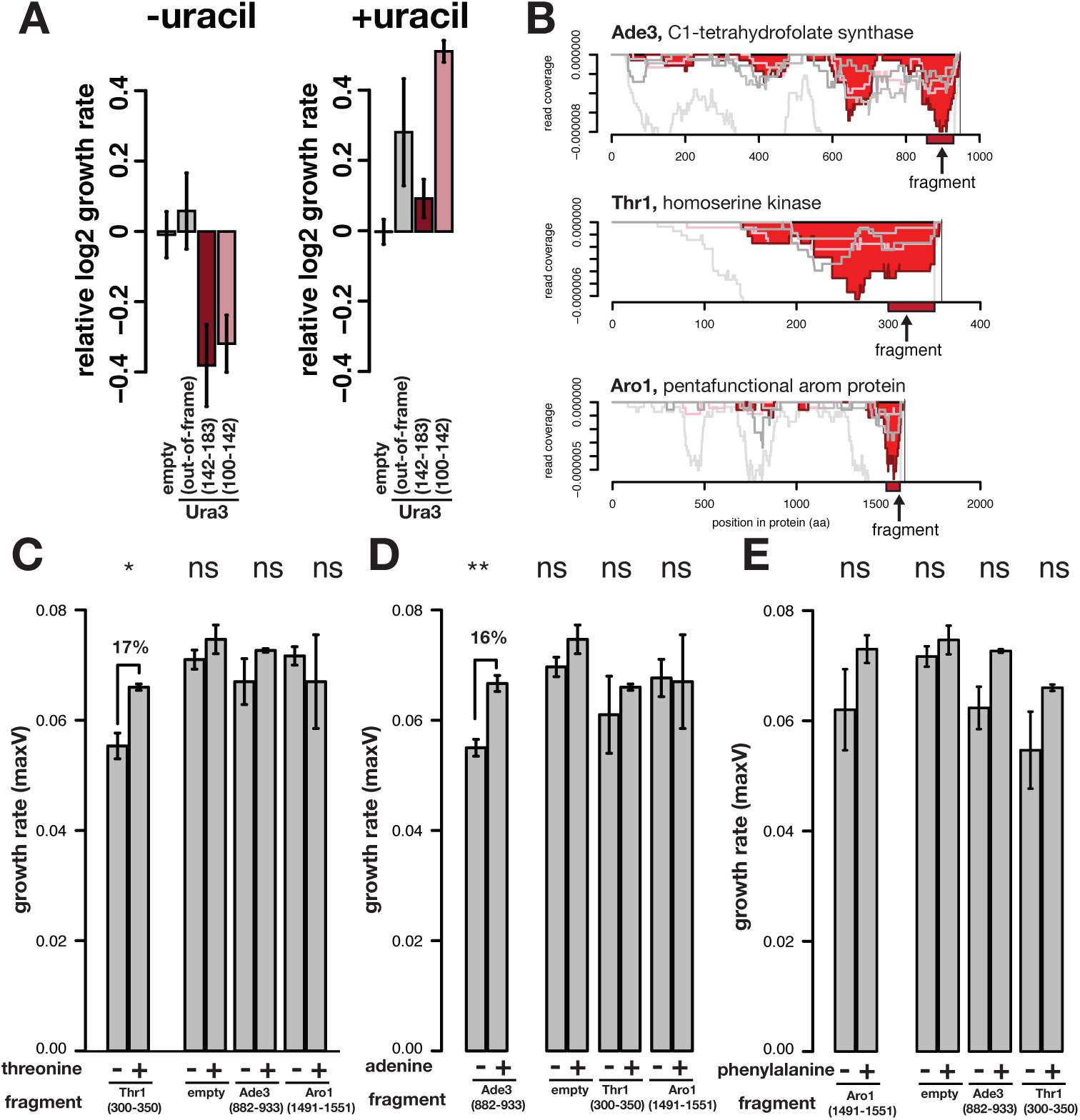
Deleterious effects of dominant negative polypeptides are exacerbated in media in which the target gene is essential. (A) Growth rates of putative dominant negative and control fragments from Ura3 selection experiment. Left panel shows growth effects in the Ura3-dependent condition of media lacking uracil (reproduced from Fig. 1D for clarity), while the right panel shows growth effects of the same fragments in the Ura3-independent condition of media containing uracil. (B) Coverage charts from the cDNA library selection in minimal media are shown for three genes related to amino acid metabolism. Our analysis identified each of these genes as having dominant negative regions. A representative fragment (indicated as red rectangles below coverage charts) was selected for each gene, and the polypeptide fragment was overexpressed in yeast cells. ¬(C-E) Barplots showing growth rates of cells containing each dominant negative fragment grown in three types of media, each lacking an amino acid related to the expected function of the target protein: (C) threonine; (D) adenine; and (E) phenylalanine. The leftmost pair of bars corresponds to the protein fragment whose function corresponds amino acid left out of the media (eg. a fragment from Thr1 is the first bar shown in (C)). For each media type, fragments not expected to affect growth were tested alongside an empty vector as control. Error bars are standard error of the mean; significance from two-sided t-test indicated with asterisks, (ns = not significant).

**Suppl. Fig. 7.**
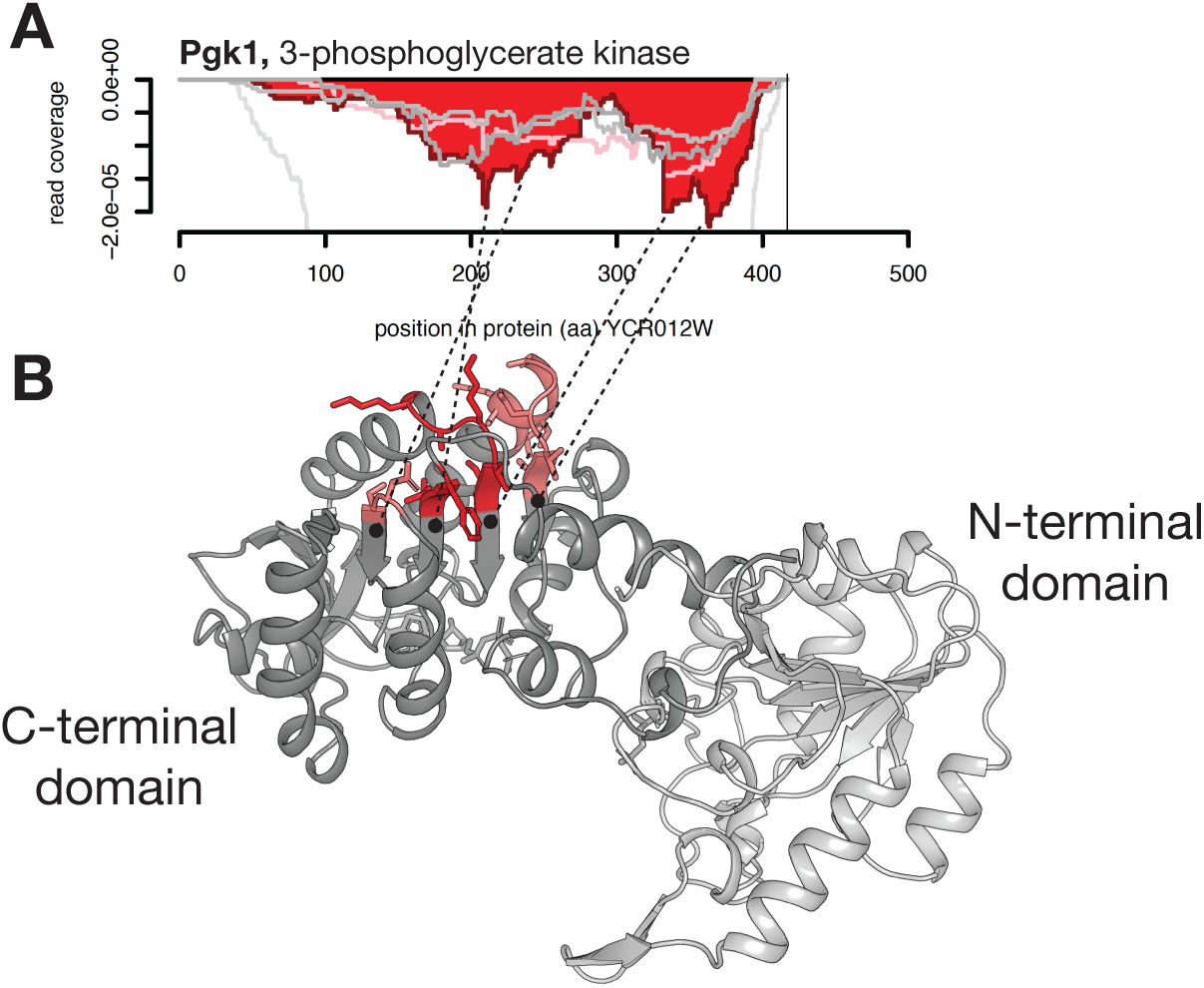
Structural features of dominant negative fragments contributing to inhibitory activity of yeast phosphoglycerate kinase. (A) Coverage chart for yeast Pgk1 showing two dominant negative regions as well as several local minima within these regions. (B) Crystal structure of the yeast Pgk1 protein, with N-terminal domain shown in light gray, and C-terminal domain shown in dark gray. Short segments colored in red correspond to the local minima in the coverage chart (dashed lines); these are found in close physical proximity and in similar locations within four β-strands.

